# Involvement of translocon complex in hemoglobin import from infected erythrocyte cytoplasm into the *Plasmodium* parasite

**DOI:** 10.1101/2021.08.02.454712

**Authors:** Priya Gupta, Rajan Pandey, Vandana Thakur, Sadaf Parveen, Inderjeet Kaur, Ashutosh Panda, Rashmita Bishi, Sonali Mehrotra, Dinesh Gupta, Asif Mohmmed, Pawan Malhotra

## Abstract

Haemoglobin degradation is crucial for the growth and survival of *Plasmodium falciparum* in human erythrocytes. Although the process of Hb degradation has been studied in great detail, the mechanisms of Hb uptake remain ambiguous to date. Here, we characterized Heme Detoxification Protein (*Pf*HDP), a protein localized in the parasitophorous vacuole, parasite food vacuole and infected erythrocyte cytosol for its role in Hb uptake. Immunoprecipitation of *Pf*HDP-GFP fusion protein from a transgenic line using anti-GFP antibody and of *Plasmodium* parasite extract using anti-human Hb antibodies respectively, showed the association of *Pf*HDP/Hb with each other as well as with the members of PTEX translocon complex. Some of these associations such as *Pf*HDP/Hb and *Pf*HDP/*Pf*exp-2 interactions were confirmed by *in vitro* protein-protein interaction tools. To know the roles of *Pf*HDP and translocon complex in Hb import into the parasites, we next studied the Hb uptake by the parasite in *Pf*HDP knock-down line using the GlmS ribozyme strategy. *Pf*HDP knock-down significantly reduced the Hb uptake in these parasites in comparison to the wild type parasites. Further, the transient knock-down of one of the members of the translocon complex; *Pf*HSP101 showed considerable reduction in Hb uptake. Morphological analysis of *Pf*HDP-HA-GlmS transgenic parasites in the presence of GlcN showed food vacuole abnormalities and parasite stress, thereby causing a growth defect in the development of these parasites. Together, we implicate the translocon complex in the trafficking of *Pf*HDP/Hb complex in the parasite and suggest a role for *Pf*HDP in the uptake of Hb and parasite development. The study thus reveals new insights into the function of *Pf*HDP, making it an extremely important target for developing new antimalarials.

## Introduction

*Plasmodium falciparum* is a leading cause of malaria with 219 million malaria cases and 4,35,000 deaths reported worldwide in the year 2018 (Organization 2019). Although drug therapies and vector control mechanisms for the containment of the disease have been developed, eradication of malaria has not been achieved yet (Rathore, McCutchan et al. 2005). The spread of resistance against the available antimalarials and unavailability of a highly efficacious vaccine poses a greater challenge to eradicate this dreaded disease. Early resistance against the artemisinin combination therapies, a present-day front-line therapy, had been reported in western Cambodia, later spreading across an expanding area of the Greater Mekong subregion (Woodrow and White 2017). Hence, there is an urgent need to identify new drug targets and molecules that can be targeted to develop effective vaccines to prevent the disease spread.

During the asexual stage of life cycle inside the human erythrocyte, the parasite digests Hb in a specialized organelle, the food vacuole. Hb is digested by the sequential action of a complex of proteases including plasmepsins, falcipains and falcilysin inside the food vacuole (Luker, Francis et al. 1996, Francis, Banerjee et al. 1997, Liu, Gluzman et al. 2005, Singh, Sijwali et al. 2006). These proteases cleave Hb to short peptides and finally to amino acids (Dalal and Klemba 2007). The free toxic by-product, heme, generated during the process is subsequently converted into an inert insoluble polymer; hemozoin (Ashong, Blench et al. 1989, Egan, Combrinck et al. 2002, Egan 2008). *Pf*HDP,(Heme Detoxification Protein) has been shown to be extremely potent in converting heme to hemozoin (Jani, Nagarkatti et al. 2008). *Pf*HDP, a food vacuole associated protein, possesses two heme binding sites and a Hb binding site (Gupta, Mehrotra et al. 2017) and is a part of a ∼200 kDa complex with other proteins including falcipain 2/2, Plasmepsin II, Plasmepsin IV, and histo-aspartic protease inside the food vacuole (Chugh, Sundararaman et al. 2013).

Although the mechanism of Hb degradation has been extensively studied; the mechanisms of uptake of the Hb from infected erythrocyte cytosol to the parasite remains poorly understood. Four distinct pathways have been proposed to assist in the uptake of Hb (Elliott, McIntosh et al. 2008, Lazarus, Schneider et al. 2008). The uptake begins with the folding of parasites around erythrocyte cytoplasm followed by the development of vesicles and cytostomes that continue uptake of Hb inside the parasite. After this step, phagosomes appear, which assist in the trafficking of Hb. Finally, cytostomal invaginations elongate to form tubes that connect to the digestive vacuole at one end and to the parasite surface at the other end, opening to the erythrocyte cytosol. Despite all the microscopic evidence available, the molecules and adaptors that participate in delivering Hb to the food vacuole hold ambiguity. Based on immuno-electron microscopy, *Pf*HDP was shown to be present in the vesicles that traffic Hb from the erythrocyte to the food vacuole of the parasite (Jani, Nagarkatti et al. 2008). *Pf*HDP lacks a classical N-terminal signal sequence or PEXEL motif, which usually assists in the sorting and transporting of any protein to a destined site using the translocon complex. The translocon complex consisting of PTEX150, Exportin 2, PTEX88, HSP101 and Trx2 is used by the parasite to export proteins from the parasite to the erythrocyte cytoplasm (de Koning-Ward, Gilson et al. 2009).. Interaction of *Pf*HDP and Hb paved the way for a hypothesis suggesting that *Pf*HDP might be playing some role in the uptake of Hb from the erythrocyte cytoplasm (Gupta, Mehrotra et al. 2017).

In this study, we attempt to unravel one of the pathways involved in the uptake of Hb from the erythrocyte cytosol. Downregulation of *Pf*HDP in the *Pf*HDP-HAGlmS transgenic parasites led the parasites to take up less Hb from the erythrocyte and induced parasite stress. Immunoprecipitation of parasite cell lysates using anti-Hb and anti-GFP antibodies showed an association of *Pf*HDP with Hb and with the components of the translocon complex, such as exportin-2, PTEX150 and HSP101. *In silico and in* vitro protein-protein interaction studies confirmed the association of *Pf*HDP with *Pf*exp-2. Hence, we propose that *Pf*HDP participates in the uptake of Hb from erythrocyte using the translocon complex. Functional insights into the role of *Pf*HDP can help us design better inhibitors targeting both the heme and Hb binding *Pf*HDP domains.

## Methods

### Maintenance of *P. falciparum* cultures and transfection

The *P. falciparum* parasite line 3D7 was maintained as described previously (Trager and Jensen 1976). To generate a GFP overexpressing transfection vector construct, the entire open reading frame of *Pf*HDP was amplified using HDPGFP-FP 5’GCAGATCTTTTTTCATCAGTATGAAAAAT AGATTTTATTAT 3’ and HDPGFP-RP 5’GC CCTAGGAAAAATGATGGGCTTATCTACTATAT3’ primer set and cloned into the pSSPF2 vector to create a C terminal *Pf*HDP-GFP fusion protein under the control of *hsp86* promoter. *P. falciparum* 3D7 ring-stage parasites were transfected with 100μg of plasmid DNA by electroporation (310 V, 950 μF) and the transfected parasites were selected using 2.5 nM blasticidin (Crabb, Rug et al. 2004). Expression of the *Pf*HDP-GFP fusion protein in transgenic *P. falciparum* blood-stage parasites was examined by western blotting and immunofluorescence. Protein bands were visualized using an ECL kit (Thermo Scientific, USA).

For transfection of knock-down constructs, C-terminal region of *Pf*HDP was amplified using gene-specific primers, HDP GlmSHA FP 5’GCAGATCTTTGAACATAAGCCTGTAAAAAGGA C 3’ and HDP GlmSHA RP 5’ GCCTGCAGAAA AATGATGGGCTTATCTACTAT3’ and cloned into the transfection vector pHA-glmS using *Pst*I and *Bgl*II restriction sites to create a fusion of desired gene of interest (GOI) with HA-glmS at the 3’ UTR under the control of native promoter. The ring-stage parasites were transfected as mentioned and transgenic parasites were selected on alternate WR22910 drug ON and OFF cycles to ensure genomic integration of *Pf*GOI-HA-glmS constructs. The transgenic parasites were then subjected to clonal selection by serial dilution to obtain parasite line from a single genome integrated clone. The integration was checked by PCR amplification of genomic fragments using different sets of primers: (1) (a/b) HDP GlmSHA FP/ HDP GlmSHA RP (2) (c/d) HDPint check 5’ GTAGAATGTATTTTTCATCAGT3’/ HAintcheck 5’TACGGATACGCATAATCGG3’.

### Cloning, Recombinant Expression, and Purification of exportin 2 protein

The forward and reverse primers used for the cloning of C-terminal *Plasmodium falciparum* exportin 2 (*Pf*exp-2) were, 5’ CCCCATGGGATCCATGAACAATTAAAGATATTTA 3’ and 5’ GCGCGGCCGCTTCTTTATTTTCATCTTTTTT 3’, respectively. The PCR product of these primers was cloned into a pJET vector and subsequently subcloned into a pET28b vector. The gene cloned in the pET-28b vector was expressed in codon+ *Escherichia coli* cells and protein localized to inclusion bodies, which were isolated as described. Briefly, the inclusion bodies were solubilized in an 8 M urea buffer (500 mM Tris, 150 mM NaCl). The suspension was incubated for 1 h at room temperature (RT) and then centrifuged at 12,000 × *g* for 30 min at RT. The supernatant containing solubilized protein was kept for binding with Ni-NTA^+^ resin overnight at RT with constant shaking. After binding, the suspension was packed in a purification column, and flow-through was collected. The resin was washed with an 8 M urea buffer containing 10 mM imidazole. Bound protein(s) was eluted in an 8 M urea buffer containing different concentrations of imidazole. Eluted protein fractions were analyzed by 12% SDS-PAGE. The eluted fractions containing the purified protein, were pooled and concentrated. The refolding method was adopted from a standard universal protocol (Tsumoto, Ejima et al. 2003). The protein was refolded gradually by decreasing the urea concentration (6M, 4M, 2M, 1M, 0M) in the refolding buffer (0.05 M Tris, pH 8, 1 mM EDTA, 0.5 M arginine, 0.4 mM Triton X-100, 1 mM reduced glutathione, 0.5 mM oxidized glutathione). The refolded proteins were concentrated and dialyzed against 0.05 M Tris, pH 8, and 0.15 M NaCl and stored at −80°C. The purified protein was analyzed on a 12% SDS polyacrylamide gel followed by western blotting with anti His HRP antibody.

### Generation of antibodies against recombinant *Pf*HDP and *Pf*exp-2

Antibodies against recombinant *Pf*HDP and *Pf*exp-2 were raised in mice and rabbit. For this 5-6-week-old female BALB/c mice were immunized with 25 µg of recombinant HDP protein emulsified in Freund’s complete adjuvant on day 0 followed by three boosts of proteins emulsified with Freund’s incomplete adjuvant on days 14, 28 and 42. The animals were bled for serum collection on day 49. In case of rabbits, NewZealand white female rabbits were immunized with 200µg of recombinant *Pf*HDP and *Pf*exp-2, respectively, emulsified in Freund’s complete adjuvant on day 0 followed by three boosts emulsified with Freund’s incomplete adjuvant on days 21, 42 and 63. The animals were bled for serum collection on day 70. The antibody titer in serum samples were quantified by enzyme-linked immunosorbent assay (ELISA).

### *In vitro* protein-protein interaction analysis

ELISA based protein-protein interaction analysis was performed as described previously (Paul, Deshmukh et al. 2017). Briefly, a 96-well microtiter plate was coated overnight at 4°C with 50 ng recombinant *Pf*HDP protein. Another unrelated *Plasmodium falciparum* recombinant protein, Ddi was coated as negative control. After blocking the wells with 5% milk in PBS for 2h, recombinant *Pf*exp-2 was added in different amounts ranging from 0 to 100ng, and the plate incubated for 3 h at 37°C. The interaction was detected using antibodies against exportin 2 (1:500). Incubation with HRP conjugated anti-rabbit antibodies (1:3000) was done for 1h and quantified after adding the substrate OPD by measuring the resulting absorbance at 490 nm.

Far western assays were performed according to the protocol described earlier (Wu, Li et al. 2007). Briefly 1-5 µg of recombinant *Pf*HDP and an unrelated *Plasmodium* protein *Pf*MLH/MBP were separated by SDS-PAGE and transferred to a nitrocellulose membrane. The proteins were first denatured and then renatured on the membrane itself. The membranes were blocked with 5% skimmed milk and incubated with 2 µg/mL of purified interacting bait proteins i.e, Hb and exp 2 in protein binding buffer (100 mM NaCl, 20 mM Tris (pH 7.6), 0.5 mM EDTA, 10% glycerol, and 1 mM DTT) for 2h at room temp. After washing the non-specific proteins, membranes were incubated with rabbit anti-Hb/ anti-exp-2 followed by incubation with HRP conjugated anti-rabbit antibodies (1:100000) for 1h at RT. Finally, membranes were imaged with a Biorad ECL chemidoc.

For *in vitro* co-immunoprecipitation, 1μg of each protein *Pf*HDP and Hb were incubated together at room temperature for 2h in a reaction volume of 100μL containing 1× binding buffer (50 mM phosphate buffer at pH 7.0, 75 mM NaCl, 2.5 mM EDTA at pH 8.0, and 5 mM MgCl_2_), 0.1% Nonidet P 40, and 10 mM DTT. The reaction mix was incubated for 2h at 4°C with 20μL of pre-equilibrated Protein A/G conjugated antibody beads. The beads were centrifuged at 1,000× g for 5 min, washed with 200μL of binding buffer containing 400 mM NaCl, boiled for 5 min in SDS/PAGE reducing loading buffer, electrophoresed, immunoblotted, and probed. Protein A/G beads conjugated to preimmune sera were used as a negative control.

The Surface Plasmon Resonance analysis was carried out on the Biacore T200 instrument (GE Healthcare). Over 9500 Response Units of the *recombinant* HDP protein were immobilized on S-Series CM5 sensor chip (GE Healthcare) using 10mM sodium acetate pH4.0 solution (GE Healthcare). The surface of the sensor chip was blocked with 1M ethanolamine-HCl pH8.5 (GE Healthcare). Recombinant *Pf*exp-2/Hb at increasing concentrations was injected over the immobilized HDP and on the reference flow cell at a flow rate of 20 μl min−1. The kinetic parameters of the interaction and binding responses in the steady-state region of the sensogram were analyzed using Biacore evaluation software, version 4.1.1 (GE Healthcare).

### Indirect immunofluorescence assay

Briefly, thin smears of parasite cultures were made on a glass slide and fixed with a mixture of methanol/acetone. Slides were blocked in blocking buffer (PBS, 3% BSA) for 2h at 37 °C. Immunostaining was performed using primary antibodies (anti-*Pf*HDP antibody 1:100, anti-Hb antibody 1:100, anti-*Pf*exp-2 antibody 1:50, anti-*Pf*PTEX150 antibody 1:50) and appropriate secondary antibody Alexa flour 594 goat anti-mice (1:500) and Alexa flour 488 goat anti-rabbit (1:500). For liquid staining the parasite samples were fixed in 4% paraformaldehyde/ glutaraldehyde. The fixed samples were permeabilized using 0.1% triton X100. The cells were blocked in 10% FBS for 2h at RT. Immunostaining was then performed using primary antibodies overnight at 4°C. Appropriate secondary antibody, Alexa flour 594 goat anti-mice (1:500) and Alexa flour 488 goat anti-rabbit (1:500) was then added to stain the parasites for 1h at RT. The nucleus of the parasites was stained using DAPI. For imaging, a drop of the suspension was taken on a slide and viewed under the microscope.

The transgenic parasite suspension was incubated with DAPI (2 ng/ml) in PBS at RT for 10 min and parasites were observed under a microscope to visualize the GFP expression. The images were captured using a Nikon A1 Confocal Microscope and exported as 8-bit RGB files. Images were analyzed using Nikon NIS Elements v 4.0software. Imaris image was created using the software IMARIS v 4.

### Immunoprecipitation reaction

Immunoprecipitation experiments were performed using the Pierce Crosslink Immunoprecipitation Kit (Product #26147). Briefly, synchronized *Plasmodium falciparum* 3D7 mid-late trophozoites were enriched from the uninfected population using density-based percoll treatment. The parasite pellet obtained was treated with Streptolysin O to lyse the erythrocyte membrane. The pellet containing the parasite surrounded by the parasitophorous vacuolar membrane was washed with PBS until lysis of RBC stopped by centrifugation at 15000×g for 1 min and the pellet was then resuspended in PBS. The parasite pellet was then lysed using the RIPA buffer (250mM Tris, 150mM NaCl, 1mM EDTA, 1% NP-40, 5% glycerol: pH 7.4) containing protease and phosphatase inhibitor cocktails (Roche) for 30 min at 4°C with intermittent mixing. Lysate was clarified by centrifugation at 15000×g for 30 min. The supernatant protein concentration was determined by the BCA Protein estimation assay kit (Pierce) using BSA standards as reference. Approximately 1mg of total protein was incubated with about 10μg of anti-Hb antibody, cross-linked to 10μl of Protein A agarose beads using disuccinimidyl suberate (DSS) as crosslinker, for 12h at 4°C with constant mixing. An equal amount of protein was allowed to bind to beads conjugated to the preimmune antibody as a control. Following binding, beads were washed with the wash Buffer and the bound proteins were eluted from the beads using the Elution Buffer (Tris-Glycine pH 2.8). Proteins in the immunoprecipitated samples were digested by in-solution trypsin digestion. Samples were brought to a final volume of 100μl in 50mM Ammonium Bicarbonate (Sigma, U.S.A.) buffer to adjust the pH to 7.8, reduced with 10mM DTT for 1h at RT followed by alkylation with 40 mM Iodoacetamide (Sigma, U.S.A.) for 1h at RT under dark conditions. Proteins were digested by the addition of Promega sequencing grade modified trypsin (V511A) at a ratio 1:50 (w/w) of trypsin: protein. For complete digestion, samples were placed in a water-bath at 37°C for 16h. After digestion, extracted peptides were acidified with 0.1% formic Acid and analyzed by mass spectrometry.

The SLO-treated trophozoite stage lysate of *Pf*HDP-GFP transgenic parasites was immunoprecipitated using GFP-Trap^®^_A Kit (Chromotek)/ anti-GFP antibody following the manufacturer’s instructions. GFP-Trap^®^_A beads/ anti-GFP antibody was allowed to bind to parasite lysate by tumbling the tube end-over-end. Proteins were eluted in a 50μl elution buffer, digested with trypsin and peptides were analysed by mass spectrometry. 3D7 parasites treated using the same protocol was used as negative control.

### Conditional knock down assay

The functional role of *Pf*HDP was determined by knocking down the HDP mRNA with glucosamine. Effect of knock down on parasite invasion was evaluated with 3D7 strain of *P. falciparum* as the control. The parasite lines (*Pf*HDP-HA-GlmS transgenic and 3D7) were synchronized using 5% sorbitol and the growth assay was set at the mid ring stage with a haematocrit and parasitemia of synchronized ring stage culture adjusted to 2% and 1%, respectively. Glucosamine was added to the parasite culture at varying concentrations (0, 1.25mM, 2.5mM, and 5mM). Parasite growth was monitored microscopically by Giemsa-stained smears. The parasitemia was estimated after an incubation of 40h in the next cycle and also in the second cycle using flow cytometry. Briefly, cells from samples were pelleted and washed with PBS followed by staining with ethidium bromide (10μg/ml) for 20min at 37°C in the dark. The cells were subsequently washed twice with PBS and analyzed on FACS calibur (Becton Dickinson) using the Cell Quest software. Fluorescence signal (FL2) was detected with the 590nm band pass filter using an excitation laser of 488nm collecting 100000 cells per sample. Uninfected RBCs stained in similar manner were used as control. Following acquisition, data were analyzed for percentage parasitemia of each sample by determining the proportion of FL2-positive cells using Cell Quest.

### Protein-protein docking analysis

PlasmoDB release (release 48) was used to retrieve sequences of *Pf*exp-2 and *Pf*HDP proteins (Bahl, Brunk et al. 2002). The predicted 3D model of *Pf*HDP was used as reported previously (Gupta, Mehrotra et al. 2017). The cryo-EM structure of *Pf*exp-2 has been resolved (Ho, Beck et al. 2018), hence its corresponding PDB structure (6E10.pdb) was retrieved from the RCSB PDB. The protein-protein docking was performed using PatchDock based on shape complementarity principles (Schneidman-Duhovny, Inbar et al. 2005). Energy refinement was performed for top 100 docked conformations using FireDock (Mashiach, Schneidman-Duhovny et al. 2008). Different conformations showing global energy ≤ -5.0 kcal/mol for *Pf*exp-2-*Pf*HDP complexes were further analysed. PyMol was used to visualize protein complexes and generate images (https://pymol.org/2/). DIMPLOT was used to retrieve and visualize residues within 4Å of interacting docked *Pf*exp-2-*Pf*HDP complexes (Laskowski and Swindells 2011). Protein Interactions Calculator (PIC) was used to identify *Pf*exp-2-*Pf*HDP interactions which recognizes various kinds of interactions, such as disulphide bonds, hydrophobic interactions, ionic interactions, hydrogen bonds, aromatic-aromatic interactions, aromatic-sulphur interactions, and cation – π interactions within a protein or between proteins in a complex (Tina, Bhadra et al. 2007).

## Results

### *Pf*HDP interacts with Hb as well with the members of *Plasmodium* translocon complex

To identify the protein(s) associated with *Pf*HDP, we generated a parasite line expressing C-terminal GFP-tagged HDP in *Plasmodium falciparum*. *Pf*HDP-GFP was expressed in an episomal construct; pSSPF2 vector (Sup Fig. 1A-D) with GFP tag at 3’ end under the control of HSP86 promoter. Western blot analysis using anti-GFP antibodies recognised two bands in *Pf*HDP-GFP in transgenic parasite lysates, one corresponding to *Pf*HDP-GFP ∼50kDa and the other at ∼25kDa. The lower band could be a result of processing of HDP-GFP in the lysate samples (Sup Fig. 1E). Indirect immunofluorescence assay of transgenic *Pf*HDP-GFP asexual blood stages showed that *Pf*HDP is localized in the vesicles transported to the food vacuole (Fig. 1A). Co-staining of these transgenic parasites with BODIPY ceramide stain followed by live cell imaging revealed that *Pf*HDP-GFP vesicles are present at the parasitophorous vacuole membrane as well as near the food vacuole indicating that *Pf*HDP is trafficked to the food vacuole as well as to the erythrocyte cytoplasm (Fig. 1B). To ascertain whether *Pf*HDP-GFP was trafficked to the erythrocyte cytosol, we fractionated the transgenic parasites using streptolysin O to separate the parasite and erythrocyte cytoplasm fractions. Western blot analysis of both fractions using anti-GFP antiserum recognized bands at ∼50kDa and ∼25kDa in parasite cytoplasm and a band at ∼25kDa in erythrocyte cytoplasm (Sup Fig. 1F). Together, these results demonstrated that *Pf*HDP-GFP was trafficked to the erythrocyte cytosol besides being transported to the food vacuole for the hemozoin formation. Western blot analysis of Streptolysin O treated fractions of infected erythrocytes and immunofluorescence assays using anti-*Pf*HDP antibodies further validated that *Pf*HDP is present both inside the parasite as well as in the erythrocyte cytoplasm (Sup Fig. 2A-B).

**Fig. 1.**
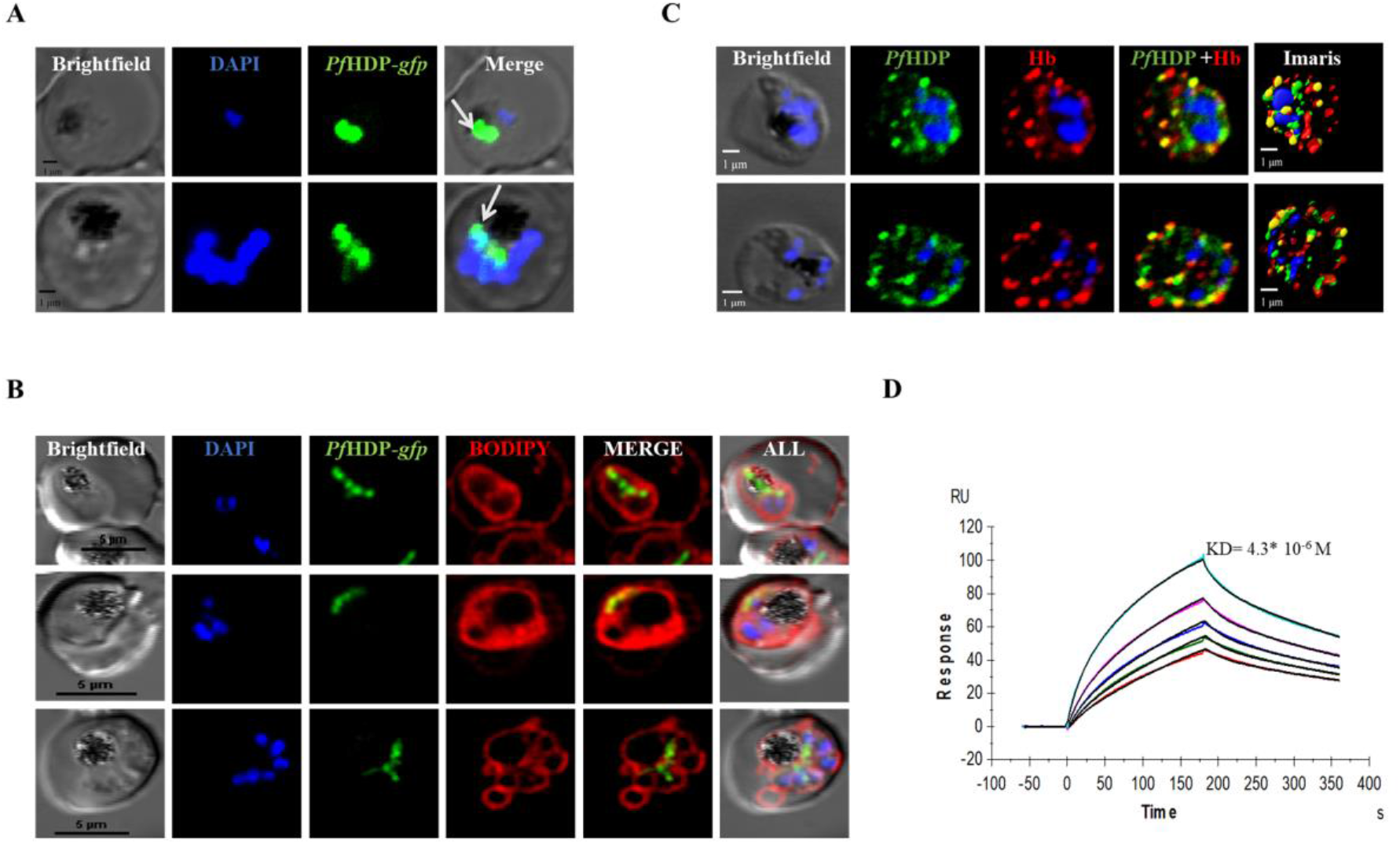
A. Subcellular localization of *Pf*HDP in *Pf*HDP-GFP transgenic lines. The image shows *Pf*HDP is trafficked to the vesicles of the parasite B. The BODIPY-TR ceramide stain, which stains the lipid membranes shows *Pf*HDP-GFP are trafficked from vesicles to the parasite plasma membrane (C) *Pf*HDP colocalizes with Hb, both inside the parasite as well as in the cytoplasm of erythrocyte (pearson coefficient co-relation - 0.8) (D) *Pf*HDP interacts with Hb in an SPR experiment. The interaction is a two-state reaction, and the observed dissociation constant is 4.3* 10^−6^ M for the reaction.

We next examined the *Pf*HDP-GFP interactome in the *Pf*HDP-GFP parasite line using a GFP pull-down assay. Briefly, *Pf*HDP-GFP protein was pulled down from cell lysates together with interacting partners, if any, using GFP-Trap beads bound with GFP antisera. Bound and eluted proteins were digested with trypsin and the released peptides were analysed by mass spectrometry to identify the interacting partners. In addition to the food vacuole proteases like Plasmepsin and falcipain 2, which are already shown to be a part of hemozoin formation complex (Chugh, Sundararaman et al. 2013), we found several proteins in the pull-down results including Hb as an interacting protein of *Pf*HDP. Additionally, members of the translocon complex including *Pf*exp-2, PTEX150, PTEX88 were identified in the immunoprecipitants (Table1). None of these proteins were pulled down from the lysates of *P. falciparum* 3D7, which served as a negative control. Together, these results suggested an association of *Pf*HDP with Hb as well as with the components of the translocon complex. To confirm the involvement of translocon complex in the trafficking of *Pf*HDP-Hb complex inside the infected asexual blood stage parasites, immunoprecipitation of Streptolysin O clarified trophozoite stage parasite extract was performed using anti-human Hb antibody with 3D7 lysate and immunoprecipitate was subjected to mass spectrometric analysis. *Pf*HDP protein was identified as an interacting partner of Hb inside the parasite along with Exp-2, PTEX150, PTEX88, HSP101 and Trx2 proteins (Table 2). None of these proteins were detected in the immunoprecipitate with pre-immune serum.

**Table 1.**
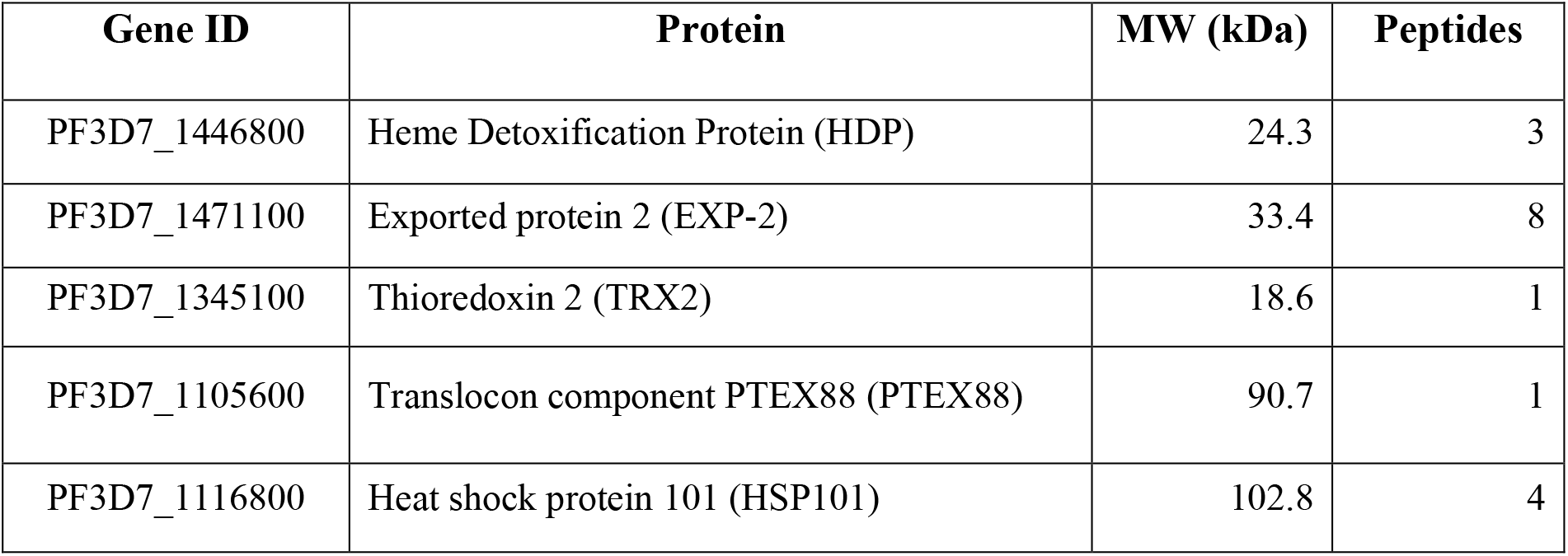
List of proteins pulled down by GFP-Trap beads from lysates of *Pf*HDP-GFP parasites from *P. falciparum*.

**Table 2.** List of proteins pulled down by anti-Hb antibody from lysates of *P. falciparum* parasites.

| Gene ID | Protein | MW (kDa) | Peptides |
| --- | --- | --- | --- |
| PF3D7_1436300 | Translocon component PTEX150 (PTEX150) | 25.74 | 7 |
| PF3D7_1105600 | Translocon component PTEX88 (PTEX88) | 90.7 | 1 |
| PF3D7_1471100 | Exported protein 2 (EXP-2) | 33.4 | 5 |
| PF3D7_1116800 | Heat shock protein 101 (HSP101) | 102.8 | 7 |
| PF3D7_1345100 | Thioredoxin-2 (TRX-2) | 18.6 | 5 |

To provide further evidence(s) for the association between Hb and *Pf*HDP, co-localization and *in vitro* protein-protein interaction studies such as co-immunoprecipitation and far-western analysis were performed. For the far-western analysis, recombinant HDP protein that served as a bait was resolved on the SDS-PAGE gel and Hb was allowed to interact with bait protein on the nitrocellulose membrane, bound protein was probed with anti-Hb antibody. A high affinity interaction between *Pf*HDP and Hb was observed (Sup Fig. 3A). Secondly, we performed interaction studies between *Pf*HDP and Hb and co-immunoprecipitation analysis was carried out using anti-HDP antiserum. As seen in Sup Fig. 3B, anti-HDP antisera could pull down Hb from a mixture of the two proteins as analyzed by western blot analysis using anti-human Hb antibody. Recombinant ClpQ protein, a mitochondrial protein, was used as a negative control. Immunolocalization studies were next performed to localize *Pf*HDP and human Hb protein together in trophozoite stage parasites using anti-*Pf*HDP and anti-Hb antibodies. *Pf*HDP co-localized considerably with human Hb with a Pearson coefficient of 0.81, both inside the parasite as well as in the erythrocyte cytosol (Fig. 1C). We further analysed the kinetics of the binding of *Pf*HDP to Hb using the Surface Plasmon Resonance. The recombinant HDP protein was immobilized on a CM5 SPR chip and Hb was allowed to interact with the immobilized protein at different concentrations ranging from 31.125µg/ml to 500µg/ml. The sensogram showed a dose dependent increase in binding of the Hb with time to the immobilized *Pf*HDP protein with an equilibrium dissociation constant of 4.3* 10^−6^ M (Fig. 1D). The binding followed a two-state reaction suggesting more than one binding site for binding of *Pf*HDP to Hb.

To validate the involvement of components of translocon complex in Hb/*Pf*HDP trafficking if any, co-localization studies were performed by immune-staining trophozoite stage parasites with the respective antibodies. Immunofluorescence assay showed that *Pf*HDP and Hb indeed co-localized with the two components of the translocon complex: *Pf*exp-2 and *Pf*PTEX150, at the parasitophorous vacuolar membrane (Fig. 2A-B and 3A). The Pearson coefficient of the correlation was found to be above ∼0.6. Together these results point towards a likely role of the translocon complex in the transport of *Pf*HDP-Hb complex from erythrocyte cytoplasm into the parasite.

**Fig. 2.**
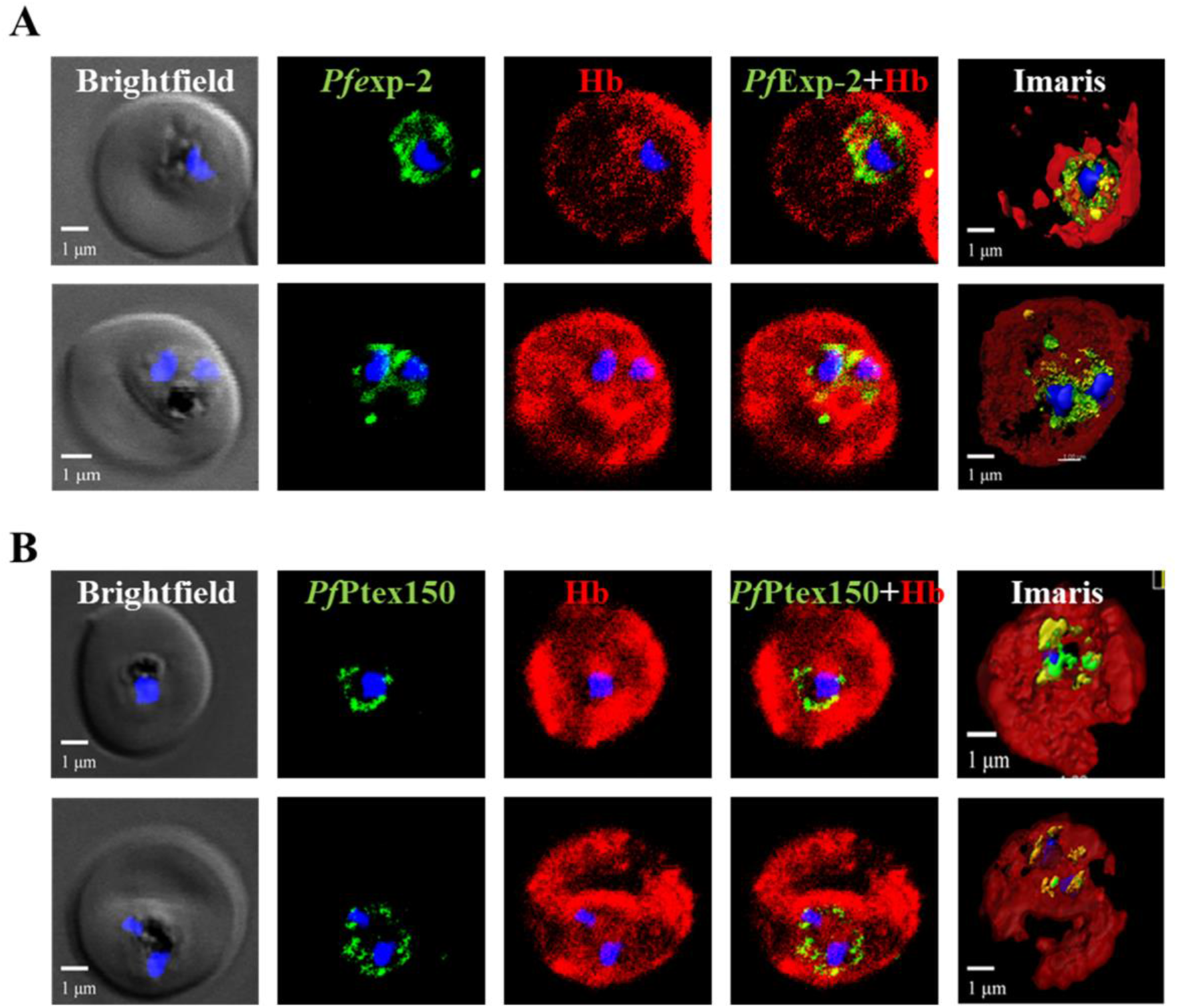
Hb colocalized with members of translocon components, *Pf*exp-2 (A) and *Pf*PTEX150 (B) with a pearson coefficient of 0.62 and 0.64, respectively.

**Fig. 3.**
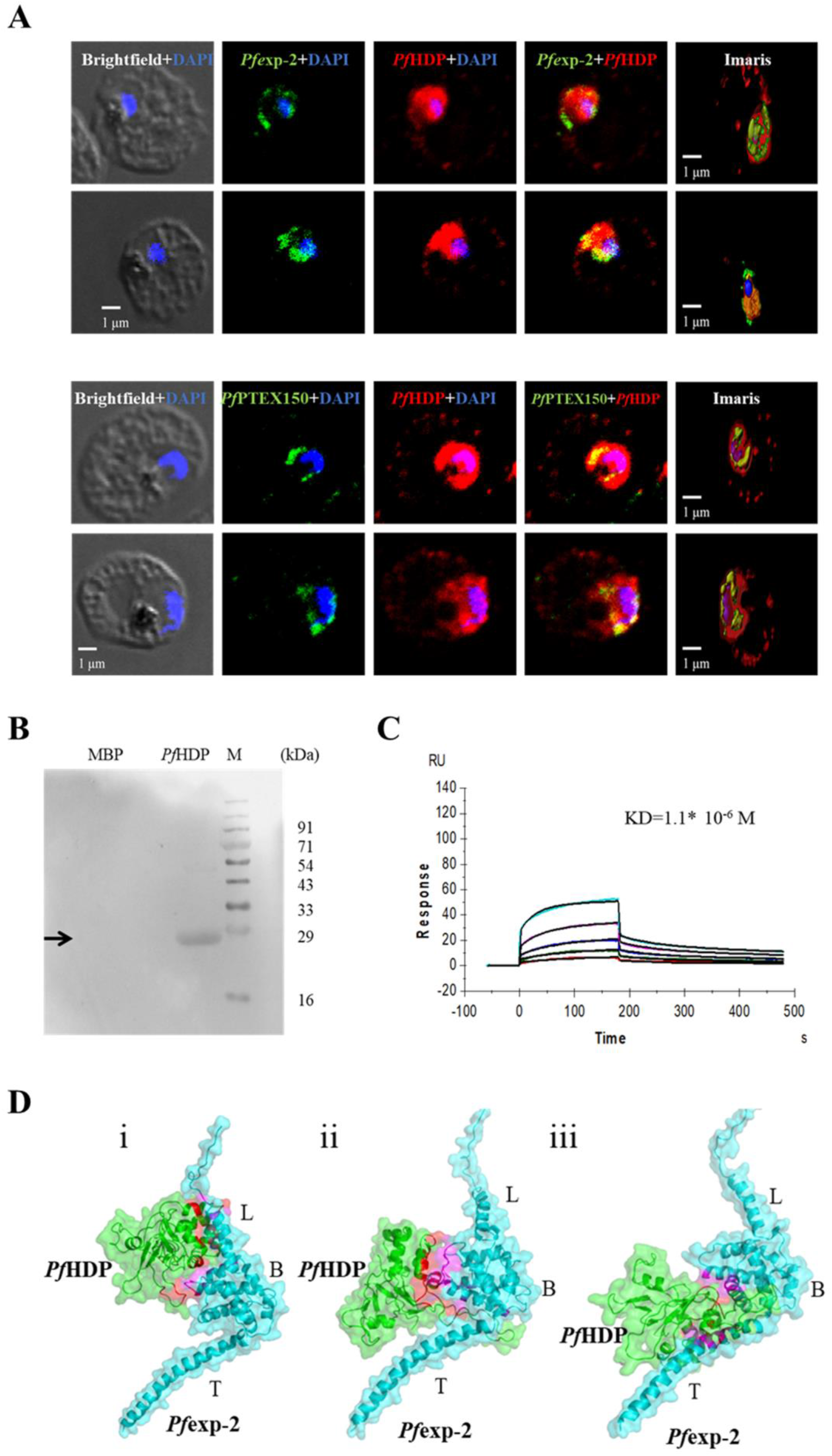
(A) *Pf*HDP colocalizes with the members of translocon components; *Pf*exp-2 and *Pf*PTEX150 at the parasitophorous vacuolar membrane with a pearson coefficient of 0.74 and 0.60, respectively. (B) *Pf*exp-2 interacts with *Pf*HDP in a far western experiment (C) *Pf*HDP interacts with recombinant C-terminal *Pf*exp-2 in an SPR experiment. The interaction is a two-state reaction, and the dissociation constant is 1.1* 10-6 M. (D) Conformational docking patterns (i-iii) observed for The *Pf*exp-2 (monomer)-*Pf*HDP complex. Green-*Pf*HDP, Cyan-*Pf*exp-2, Red – *Pf*exp-2 interacting region within 4Å of *Pf*HDP, Magenta - *Pf*HDP interacting region within 4Å of *Pf*exp-2.

### *Pf*HDP interacts with exportin 2 at the parasitophorous vacuolar membrane

We next sought to analyse the interaction of *Pf*HDP with one of the members of the translocon complex; *Pf*exp-2 that has been suggested to be a pore forming protein on the parasitophorous vacuolar membrane (Garten, Nasamu et al. 2018, Sanders, Dickerman et al. 2019). To study the interaction between *Pf*HDP and *Pf*exp-2, we cloned and expressed a C-terminal fragment of *Pf*exp-2 encompassing amino acids 139-285 in *E. coli* (Sup Fig. 4A). The protein was purified to homogeneity using the Ni-NTA^+^ column. The purified protein migrated at a molecular size of ∼20kDa as seen in Coomassie stained SDS-PAGE and western blot experiment performed using anti-His-HRP antibody (Sup Fig. 4B-C). Specific antibodies were generated against recombinant *Pf*exp-2 protein in rabbit. The antisera recognized a band of ∼33kDa in the 3D7 parasite cell lysate, which corresponds to the monomeric form of native *Pf*exp-2 (Sup Fig. 4D).

We further employed *in vitro* protein-protein interaction tools such as *in vitro* ELISA based protein binding assay and far-western binding analysis to assess *Pf*HDP interaction with *Pf*exp-2. Recombinant *Pf*HDP protein interacted with *Pf*exp-2 in a concentration dependent manner in (Sup Fig. 4E). A nonspecific recombinant *Pf*Ddi protein was used as a negative control, which showed no significant interaction with *Pf*exp-2. For the far-western binding analysis we used recombinant *Pf*HDP protein as a bait and *Pf*exp-2 was allowed to interact with bait protein on the nitrocellulose membrane and probed with anti-exp-2 antibody. A nonspecific purified MBP protein was used as a negative control. As evident in Fig. 3B, *Pf*HDP interacts specifically with *Pf*exp-2 (Fig. 3B). To know whether *Pf*HDP interacts with *Pf*exp-2 in the cell, parasite lysate from *Pf*HDP-GFP transgenic line was immunoprecipitated using GFP-TRAP column and elutes from the GFP pull down assay were analysed by western blot analysis using anti-*Pf*exp-2 antibody. The parasite lysate from 3D7 parasite line was used as a negative control. As seen in Supplementary figure 4F, *Pf*HDP-GFP fusion protein interacted with *Pf*exp-2, which was detected in the western blot analysis, whereas 3D7 lysate did not show any *Pf*exp-2 band. To study the kinetics of the binding of *Pf*HDP to *Pf*exp-2, we performed the Surface Plasmon Resonance analysis by immobilizing the recombinant HDP protein on a CM5 chip using EDC-NHS coupling. Recombinant *Pf*exp-2 was allowed to interact with the immobilized protein at different concentrations ranging from 0.625 µM to 10 µM. The sensogram showed a dose dependent increase in binding of the *Pf*exp-2 with time to the immobilized *Pf*HDP, with an equilibrium dissociation constant of 1.1* 10^−6^ M (Fig. 3C). Together, these binding studies unequivocally suggested an interaction between *Pf*HDP and *Pf*exp-2.

To identify the interaction sites for the *Pf*exp-2 and *Pf*HDP, we carried out *in silico* interaction analysis using the already known structure of the malaria translocon complex, 6E10.pdb and HDP structure model. *In silico* docking analysis of *Pf*exp-2 monomer with *Pf*HDP using PatchDock and energy refinement on top 100 conformations were carried out using default parameters. Next, the docked conformations, with global binding energy better than -5 kcal/mol were analyzed. Interestingly, the docking analysis showed that *Pf*HDP binds to the multiple sites of *Pf*exp-2, which includes Linker helix (L), globular domain body (B) and transmembrane domain (T) of *Pf*exp-2 (Fig. 3D) with highest binding energy observed for *Pf*HDP-*Pf*exp-2 in the Linker region (−39.57 kcal/mol, pose 1). Similarly, the best binding score observed for *Pf*HDP binding to the globular domain body (B) and transmembrane domain (T) of *Pf*exp-2 was -17.08 (pose 2) and -6.35 kcal/mol (pose 3), respectively. The PPI analysis showed gradual changes in the *Pf*HDP residues, which interacts with different pockets of *Pf*exp-2 (Supplementary Table 1) however *Pf*HDP TYR56 and ASN59 residues binding remain consistent and were present within 4Å of *Pf*exp-2 in almost all conformations analyzed for *Pf*exp-2-*Pf*HDP complexes (Sup Fig. 5). Thus, our bioinformatics analysis further supported an interaction between *Pf*exp-2 and *Pf*HDP.

### Knockdown of *Pf*HDP results in parasite stress and low levels of Hb inside the parasite

To illustrate the functional importance of *Pf*HDP protein in parasite biology in particularly for Hb import at asexual blood stages, a transgenic line with the genomic locus of *Pf*HDP fused to GlmS ribozyme system was generated. The GlmS ribozyme is transcribed along with the gene. This inducible knockdown system uses glucosamine as an inducer. In the presence of glucosamine, which binds to ribozyme inducing the cleavage of the chimeric mRNA and hence knocking-down the respective targets (Sup Fig. 6A). The transgenic line was generated using pHA-GlmS vector-based constructs (Sup Fig. 6B-C). The integrants were selected using WR22910 selection, followed by clonal selection by limited dilution to obtain a pure conditional knockdown parasite line. The integration was confirmed by PCR amplification using different combinations of primer sets (Sup Fig. 6D). Expression of the HA tagged fusion protein and the native protein were confirmed by western blot analysis of the transgenic parasites using anti-HA antibody, which detected a single band at ∼50kDa, corresponding to the size of dimeric native protein in the parasite (Sup Fig. 6E). No such band was detected in 3D7 parasite lysate was used as a negative control

To study the effect of the knockdown on the expression of *Pf*HDP protein, the late trophozoite stages of transgenic parasites were treated with different concentrations of GlcN (0mM, 1.25mM, 2.5mM, respectively). The parasites were harvested in the next cycle at 42-44 hpi and saponin treated parasites were lysed in RIPA buffer. Expression of the fusion protein was analysed by western blot analysis of the lysates from transgenic parasites using anti-HA antibody. A considerable reduction in *Pf*HDP protein was seen in the presence of different concentrations of GlcN (Fig. 4A). *Pf*BiP, a constitutively expressed endoplasmic reticulum chaperone protein, was used as a loading control. We next studied the effect of the knockdown of this protein on the growth of the parasites. GlcN was added at the ring stage parasites 16-20 hpi at varying concentrations (0 mM, 1.25 mM, 2.5 mM) and the growth was monitored till the formation of new rings up till two invasion cycles. In the first cycle, a slight decrease in the parasitemia was observed at 2.5mM GlcN concentration. However, when the parasites were allowed to progress to the second cycle, growth inhibition of ∼40% was observed in *Pf*HDP-HAGlmS knockdown parasites (Fig. 4B). The inability of GlcN to inhibit more than 50% parasite growth could be attributed to the incomplete knockdown of the protein in the first cycle, as the synthesis of *Pf*HDP begins as early as the ring stages of the parasite. Giemsa smears in the second cycle of growth demonstrated an induction of parasite stress and food vacuole abnormalities (Fig. 4C). The 3D7 parasites treated similarly with varied GlcN concentrations were used as a negative control. We further analysed the Hb levels in the knockdown parasites by western blotting using anti-Hb antibody. A considerable decrease in the amounts of Hb was seen inside the GlcN treated parasites (Fig. 4A). *Pf*BiP was used as a loading control and it did not show any change in expression. We next studied the expression/localization of *Pf*HDP and Plasmepsin 2 in the *Pf*HDP knock-down parasites. Interestingly, we observed an inappropriate expression for Plasmepsin 2, a food vacuole protease in these stressed parasites (Fig. 4D). Overall, these results demonstrated a role of *Pf*HDP in Hb uptake and its impact on Hz formation.

**Fig. 4.**
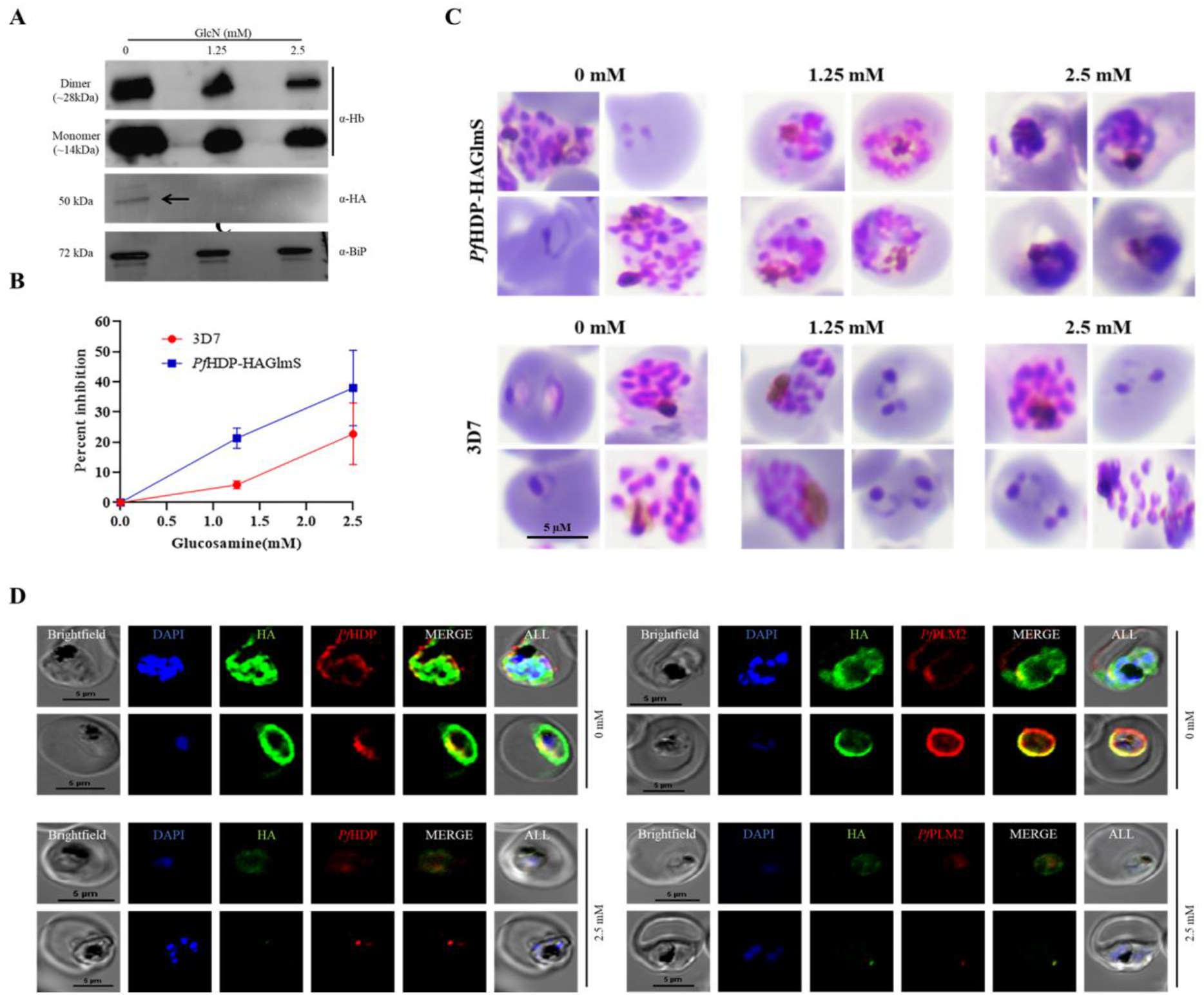
(A) Effect of conditional knockdown of *Pf*HDP in parasites showing up to 45% invasion inhibition. Data represents mean ± SD; n = 3 experiments. (B) Western blot analysis of lysate from *Pf*HDP-HA-glmS line with α-HA rat serum and α-Hb antibody. Hb uptake is reduced in the knockdown parasites. Anti BiP was used as loading control. (C) Representative Giemsa-stained smears highlighting the parasite stress in the trophozoite stage following *Pf*HDP knockdown. (D) Immunofluorescence assays show the low levels of *Pf*HDP and plasmepsin-2 in knockdown lines.

### Parasites expressing low levels of *Pf*HSP101 protein display low levels of Hb inside the parasite

To characterize the role of components of the translocon complex in Hb trafficking, in particular the Hb import, we next studied the uptake of Hb in *Pf*HSP101-DDDHA (Beck, Muralidharan et al. 2014)knockdown lines. For the same, the *Pf*HSP101-DDD-HA parasites were grown in the presence of trimethoprim (TMP) (Fig. 5A); removal of TMP causes functional interference of *Pf*HSP101 protein, thereby leading to its knockdown (Beck, Muralidharan et al. 2014). Briefly, the late trophozoite stages of parasites were treated with different concentrations of TMP (0μM, 2.5μM, 5μM, respectively) and these untreated v/s treated parasites were harvested in the next cycle at 42-44 hpi. The saponin lysed parasites were subjected to lysis in RIPA buffer and Hb levels in the control v/s knockdown parasites were analysed by western blot analysis using anti-Hb antibody. A remarkable decrease in levels of Hb was observed in the TMP untreated parasites where *Pf*HSP101 protein had been reduced considerably in comparison to the TMP treated parasites (Fig. 5B). *Pf*BiP was used as a loading control. Reduction in *Pf*HSP101 levels was confirmed in TMP untreated parasite lysates using anti-HA antibody to confirm the successful knockdown of HSP101 protein.

**Fig. 5.**
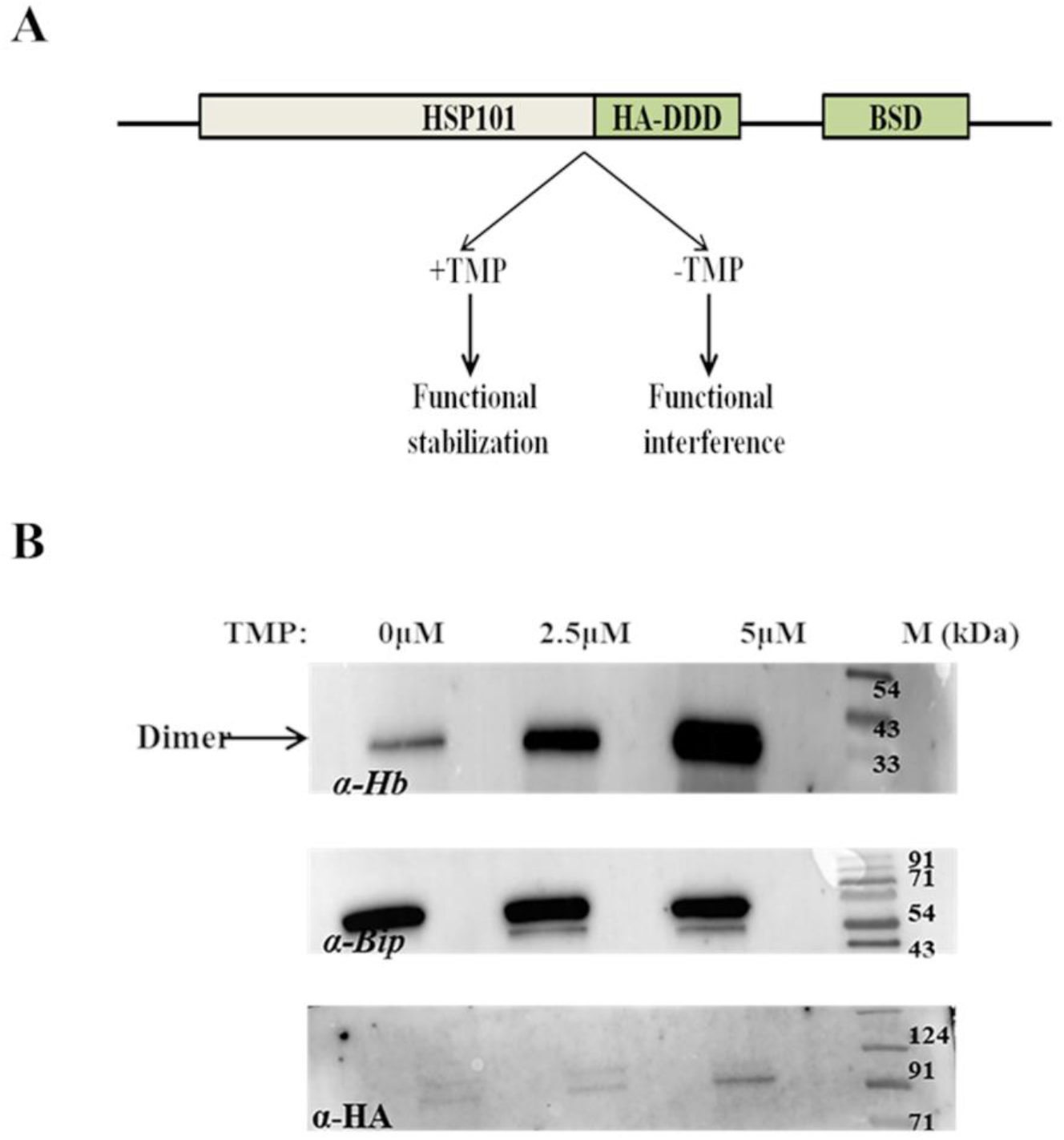
HSP101 DDDHA knockdown parasites take up less Hb from the host erythrocyte cytoplasm. (A) Illustration of the HSP101 DDDHA transgenic construct (B) western blot to detect the Hb levels inside the parasite in HSP101 knockdown parasites. BiP is used as a loading control.

In summary, these results advocated a role for *Pf*HDP in trafficking/transport of Hb into the parasite. Based on the subcellular localization of *Pf*HDP, its interaction with the Hb and with *Pf*exp-2, and its effect on the uptake of Hb in knockdown parasites, we propose a model suggesting that *Pf*HDP interacts with Hb in infected erythrocyte. The *Pf*HDP-Hb complex is then taken into the parasitophorous vacuole through the translocon complex by the interaction of *Pf*HDP with *Pf*exp-2. The *Pf*HDP-Hb complex is subsequently taken up by the parasite and gets translocated to the food vacuole by vesicular trafficking.

## Discussion

Hb uptake and its degradation are highly crucial processes for the growth of *P. falciparum*. The degradation of Hb results in the generation of amino acids that are utilized by the parasite for its protein synthesis. Heme generated as a by-product during the process of Hb catabolism is highly toxic for the survival of parasites. Heme Detoxification Protein has earlier been shown to be involved in the conversion of heme to an inert polymer hemozoin (Jani, Nagarkatti et al. 2008). In the present study, we aimed to understand Hb uptake/trafficking inside the infected parasites with a possible role of *Pf*HDP and translocon complex in Hb inward trafficking.

It has been reported earlier that *Pf*HDP is exported into the infected erythrocyte cytosol and takes a circuitous trafficking route to reach the food vacuole and catalyzes in Hz formation in the food vacuole (Jani, Nagarkatti et al. 2008). These authors further showed using anti-*Pf*HDP antibody as well as C- or N-tagged C-Myc-HDP protein that the trafficking of *Pf*HDP to the cytosol of RBCs does not occur via the classical secretory pathway and the inbound HDP protein(s) are trafficked via their cytosomal uptake and vesicular trafficking (Jani, Nagarkatti et al. 2008). However, questions that remain unanswered is that how *Pf*HDP is exported/imported into the erythrocyte cytosol or imported into the parasite back? To gain insights into the mode of trafficking of *Pf*HDP into the infected parasites from RBC cytosol, a C-terminal GFP fusion HDP overexpressing parasite line under the control of Hsp86 promoter was generated. Confocal microscopy studies showed that the inbound *Pf*HDP-GFP protein is trafficked via vesicular trafficking. SLO treatment on these transgenic parasites showed the expression of *Pf*HDP-GFP fusion protein in the SLO soup suggesting that *Pf*HDP-GFP protein is exported to PVM and to the erythrocyte cytoplasm. To further illustrate the import of *Pf*HDP, immunoprecipitation analysis of *Pf*HDP-GFP transgenic line or 3D7 line lysates with either GFP trap/anti GFP antibody or anti-Hb antibody were performed. Both the immunoprecipitates showed the presence of *Pf*HDP and Hb together along with the members of the translocon complex (de Koning-Ward, Gilson et al. 2009) such as *Pf*PTEX150, *Pf*Exp-2 and *Pf*HSP101, thereby implicating the role of *Pf*HDP and translocon complex in Hb trafficking. Earlier, we have shown that *Pf*HDP binds both heme as well as Hb (Gupta, Mehrotra et al. 2017). Based on these observations, we propose a model depicting *Pf*HDP as an adapter protein that interacts with Hb in the erythrocyte cytoplasm and helps in the intake of Hb via the translocon complex.

To shed more lights into the proposed model, we expressed a C-terminal fragment of *Pf*exp-2 and raised specific antibodies against *Pf*exp-2. Additionally, an anti-peptide PTEX-150 antibody was generated. Both, *Pf*HDP or Hb colocalized well with either *Pf*exp-2 or *Pf*PTEX150 at the parasitophorous vacuolar membrane with a pearson’s coefficient of >0.5, advocating a role for translocon complex in Hb import. A recent near atomic resolution cryoEM structure of endogenous translocon complex revealed that *Pf*exp-2 and *Pf*PTEX150 intermingle to form a static, funnel shaped pseudo-sevenfold symmetric protein conducting channel spanning the vacuole membrane (Chi-Min Ho et al., 2018). To support further on the role of *Pf*HDP protein in import of Hb from erythrocyte cytoplasm, interaction studies between *Pf*HDP and Hb or *Pf*exp-2 were performed by far western blot analysis or by Surface Plasma Resonance analysis. Recombinant HDP interacted with both Hb as well as *Pf*exp-2 with considerable affinities, thereby suggesting that *Pf*HDP-Hb complex formed in infected erythrocyte cytoplasm is translocated through the static funnel by binding to *Pf*exp-2. These results were also substantiated by docking studies between *Pf*HDP and *Pf*exp-2

To study the functional relevance of *Pf*HDP and components of translocon complex in parasite import of Hb, a *Pf*HDP-HAGlmS inducible knockdown transgenic line was generated, although our attempts to knockout *Pf*HDP gene failed. Knock-down of HDP in the parasite line using the inducer glucosamine induced food vacuole abnormalities and parasite stress. Growth inhibition was observed in glucosamine treated parasites. A reduced level of *Pf*HDP inside the transgenic lines also led to the reduction in the uptake of Hb from the parasite cytosol. These parasites also showed mis-localized or poorly expressed Plasmepsin 2 protein, a part of the hemozoin formation complex. These knocked-down parasites appeared to be in stress. The food vacuole was not properly developed in the *Pf*HDP knocked-down parasites and hence the parasites exhibited gross morphological deformities. Attempts to study the transport of Hb in *Pf*HSP101 DDDHA transgenic lines revealed that knockdown of one of the components of translocon complex reduced the ability of parasites to take up Hb from the erythrocyte cytoplasm. Hence the translocon machinery appeared to be essential for the uptake of Hb from the erythrocyte.

In summary, here we characterized *Pf*HDP for its role in Hb uptake in addition to its previously characterized function in heme degradation. To characterize the role of *Pf*HDP in Hb transport, here we generated a *Pf*HDP-GFP transgenic line. Immunoprecipitation of highly synchronized culture of trophozoites from *Pf*HDP-GFP line using anti-GFP antibody followed by LC-MS/MS analysis showed the association of Hb and *Pf*HDP with each other and with the members of translocon complex such as Exp-2, PTEX150, PTEX88, HSP101 and Trx2, a complex known to export parasite proteins. *In*silico analysis and *in*vitro protein-protein interaction techniques confirmed the interaction of *Pf*HDP with *Pf*exp-2. We further showed that *Pf*HDP is highly crucial in maintaining food vacuole and parasite health as attempts to knockdown the protein induced parasite stress. Hb uptake is severely affected in these transgenic parasites. The study thus emphasizes on the dual roles of Heme Detoxification Protein in Hb uptake as well as in conversion of heme to hemozoin. Looking at the multiple roles of *Pf*HDP in the parasite life cycle, *Pf*HDP appears to be an important target for new antimalarial discovery.

## Abbreviations

HDP: Heme Detoxification Protein
Exp-2: exportin2
Hb: hemoglobin

## Acknowledgement

This work was financially supported by Department of Biotechnology, Government of India <u>(BT/PR5267/MED/15/87/2012 and flagship project;</u> BT/IC-06/003/91) from the Department of Biotechnology, Govt. of India. PM is a recipient of the J C Bose Fellowship awarded by SERB, Govt. of India, and work is supported by the grant (DST/20/015). We thank Prof Daniel E. Goldberg for providing us the *Pf*HSP101 DDDHA transgenic parasites. We thank Dr Paul Gilson for critical review of the manuscript. We also thank the Rotary Blood Bank for providing human red blood cells for *Plasmodium* culture. P.G. was recipient of ICMR Cenetenary Post-Doctoral Fellowship, ICMR, Government of India. We thank Dr. Naresh Sahoo and Surabhi Dabral for their help in the SPR interaction experiments and confocal imaging, respectively. We thank the animal house facility for help with antibody generation in mice and rabbit.

## Ethics statement

The animal work performed in this study was approved by the Institutional Animals Ethics Committee of ICGEB (IAEC-ICGEB). Rotary blood bank provided human red blood cells.

## Author Contributions

PM conceived the idea. PM and PG designed the experiments. PG performed literature analysis. PG, RP, VT, SP, IK, AP, RB and SM performed experiments. RP and DG performed bioinformatics analysis. AM, DG and PM supervised the study. PG, PM wrote the manuscript, and all authors read and approved the manuscript.

## Supplementary Figures

**Sup fig. 1.**
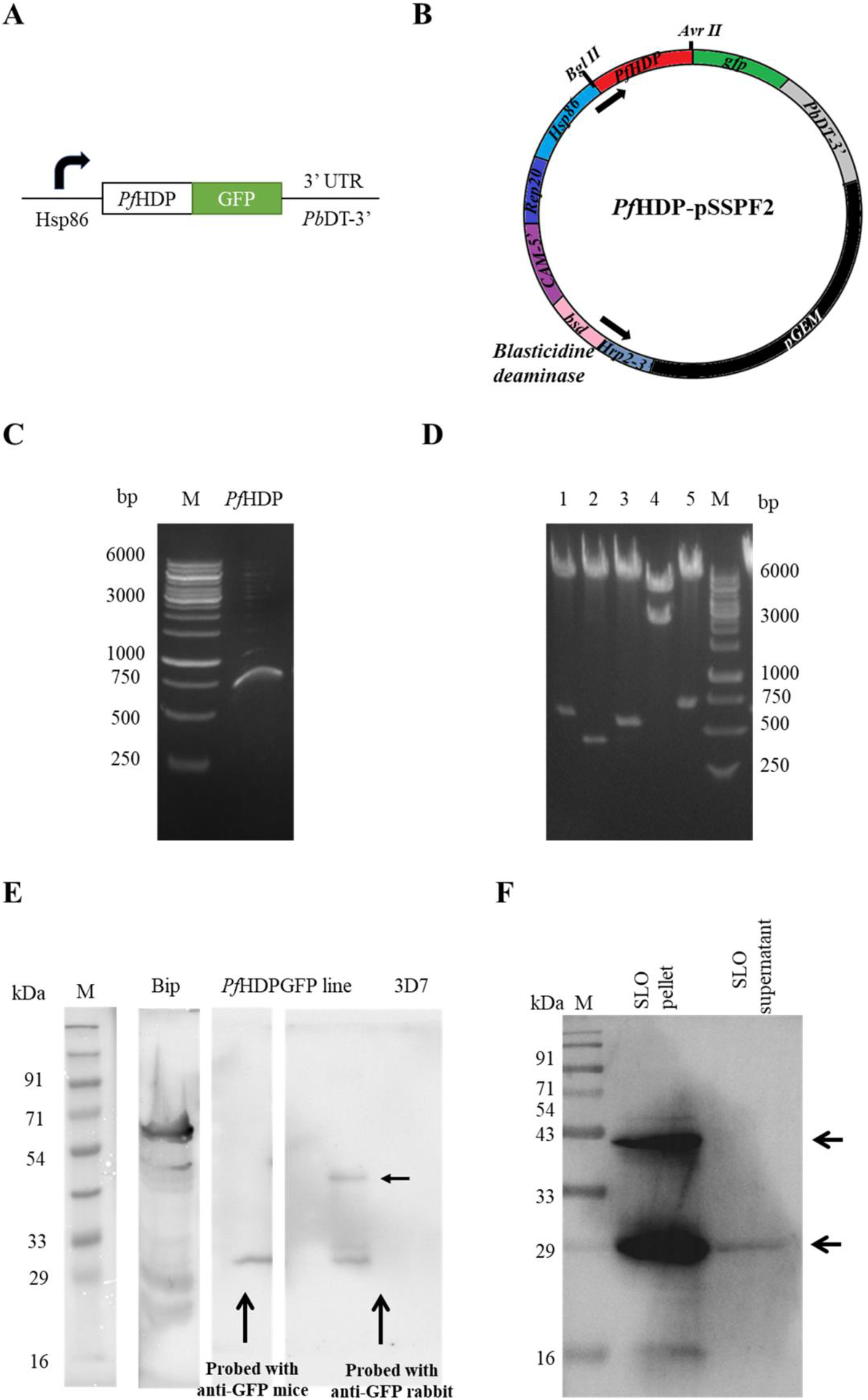
Generation of *Pf*HDP-GFP line in pSSPF2 vector. (A) Full length *Pf*HDP gene cloned in pSSPF2 vector with GFP tag at 3’ end under the control of HSP86 promoter (B) plasmid vector pSSPF2 for expressing gene products in malarial transfectants (C) PCR amplification of *Pf*HDP gene from cDNA using the *Pf*HDP-GFP FP and *Pf*HDP-GFP RP primer set. (D) The construct *Pf*HDP-pSSPF2 was checked for correct insertion of *Pf*HDP and presence of other sequence regions using different sets of restriction enzymes (Lane 1: BglII/AvrII-HDP; Lane 2: BamHI/HindIII-blasticidine resistance; Lane3: EcoRI/HindIII-HRP 2-3’, Lane 4: NotI/EcoRI-pGEM backbone, Lane 5: XhoI/AvrII: GFP (E) Western blot to confirm the expression of GFP fusion protein in the *Pf*HDP-GFP transgenic line. *Pf*BiP was used as a positive control. (F) *Pf*HDPGFP detection in the SLO pellet and supernatant fractions suggesting *Pf*HDP is transported to the erythrocyte cytosol.

**Sup fig. 2.**
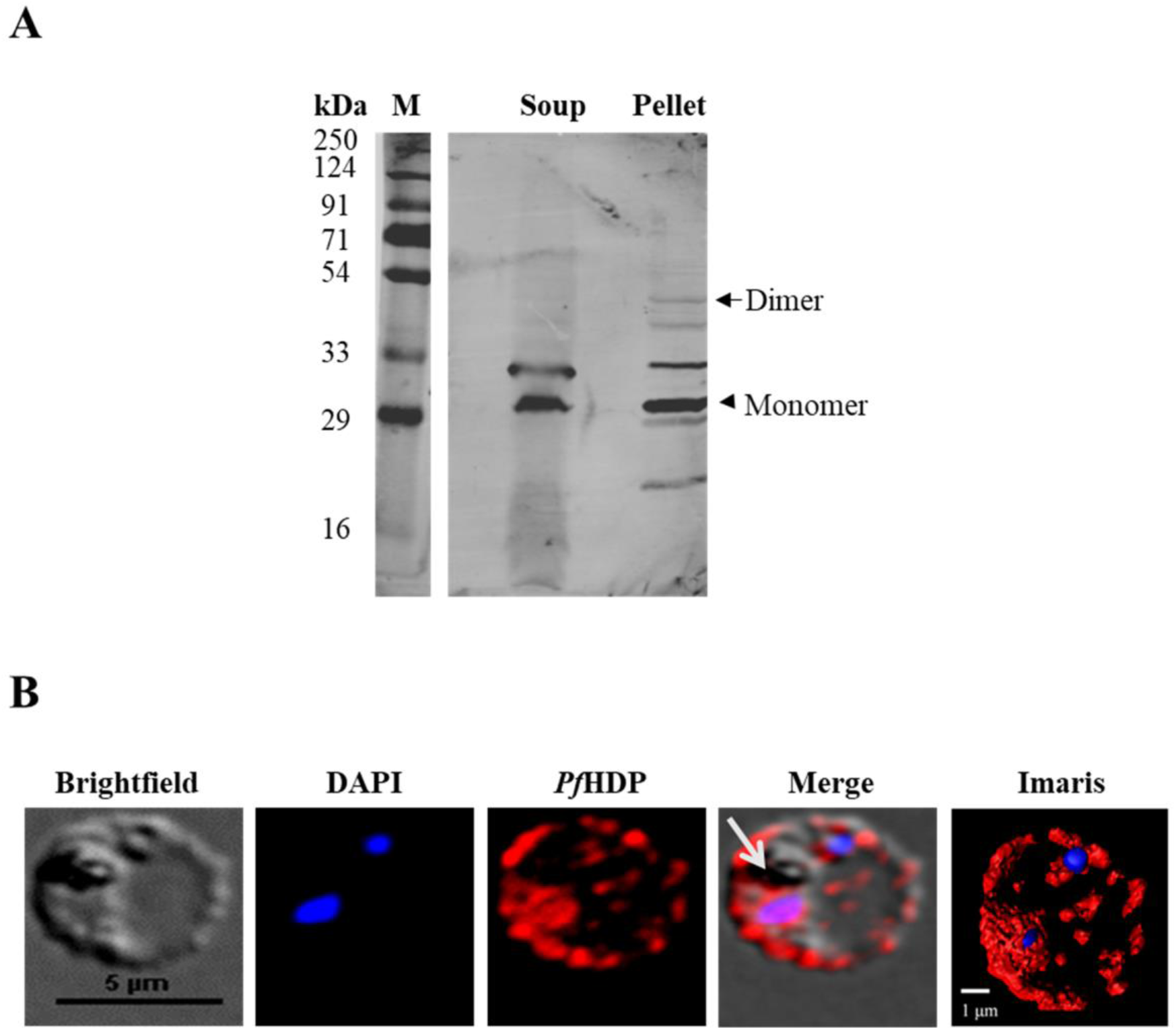
(A) *Pf*HDP is detected in both the fractions of SLO treated infected erythrocytes. The protein is detected in both the parasite cytoplasm and erythrocyte cytoplasm by *Pf*HDP antibodies in a western blot assay. (B) Immunofluorescence assay shows the localization of *Pf*HDP (red) in both the erythrocyte cytosol and parasite cytoplasm.

**Sup fig. 3.**
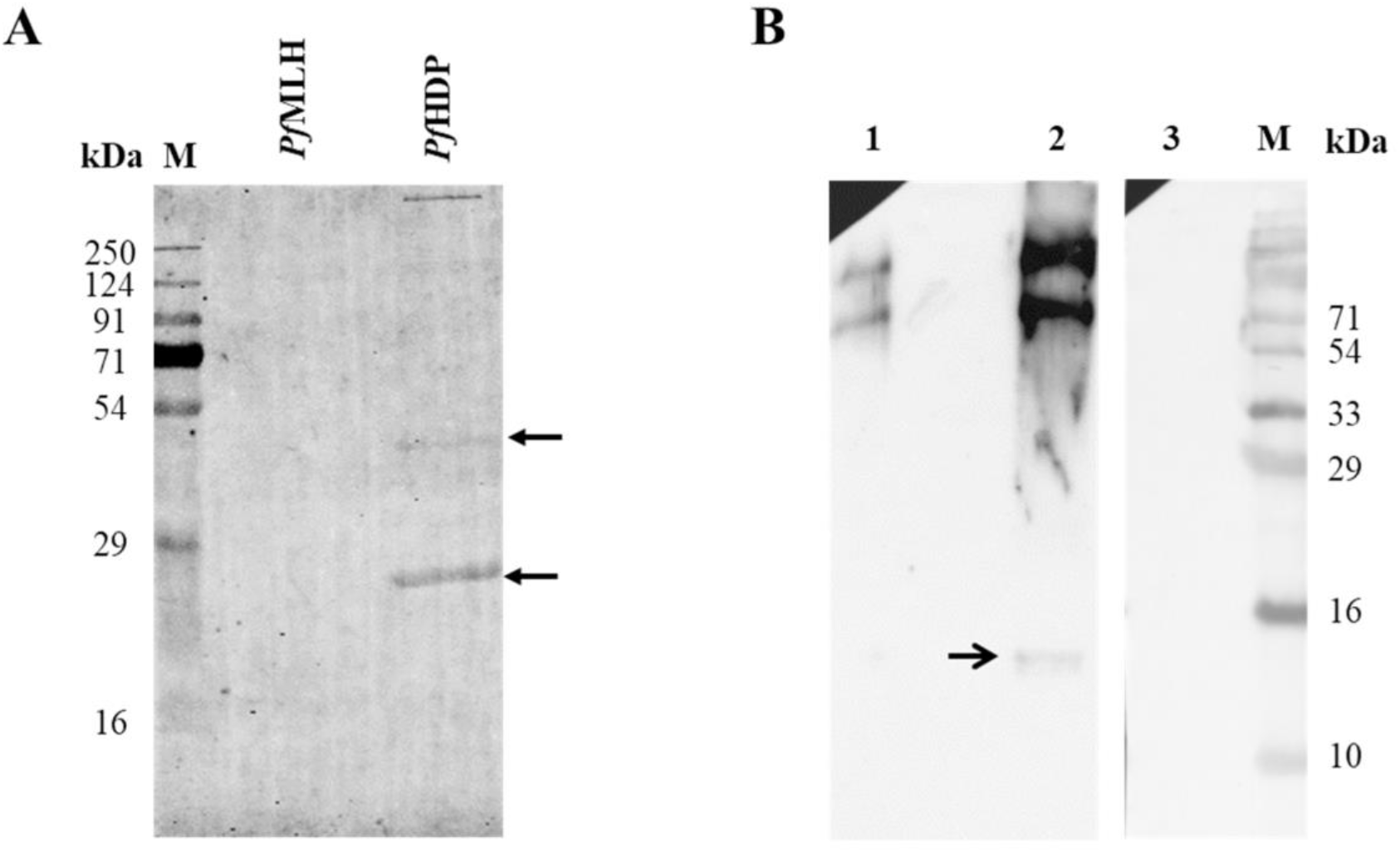
(A) *Pf*HDP interacts with Hb in a far western interaction experiment. Here, *Pf*MLH was used as a negative control. (B) *Pf*HDP and Hb interact with each other in a co-immunoprecipitation experiment. Lane 1 contains elute pulled from the mixture of *Pf*HDP and Hb using Pre immune sera. Lane 2 contains elutes pulled from a mixture of *Pf*HDP and Hb using the HDP antibody. Lane 3 contains an eluted mixture of HDP and ClpQ using the *Pf*HDP antibody. The arrow shows Hb pulled by *Pf*HDP antibody as probed by Hb antibody.

**Sup fig. 4.**
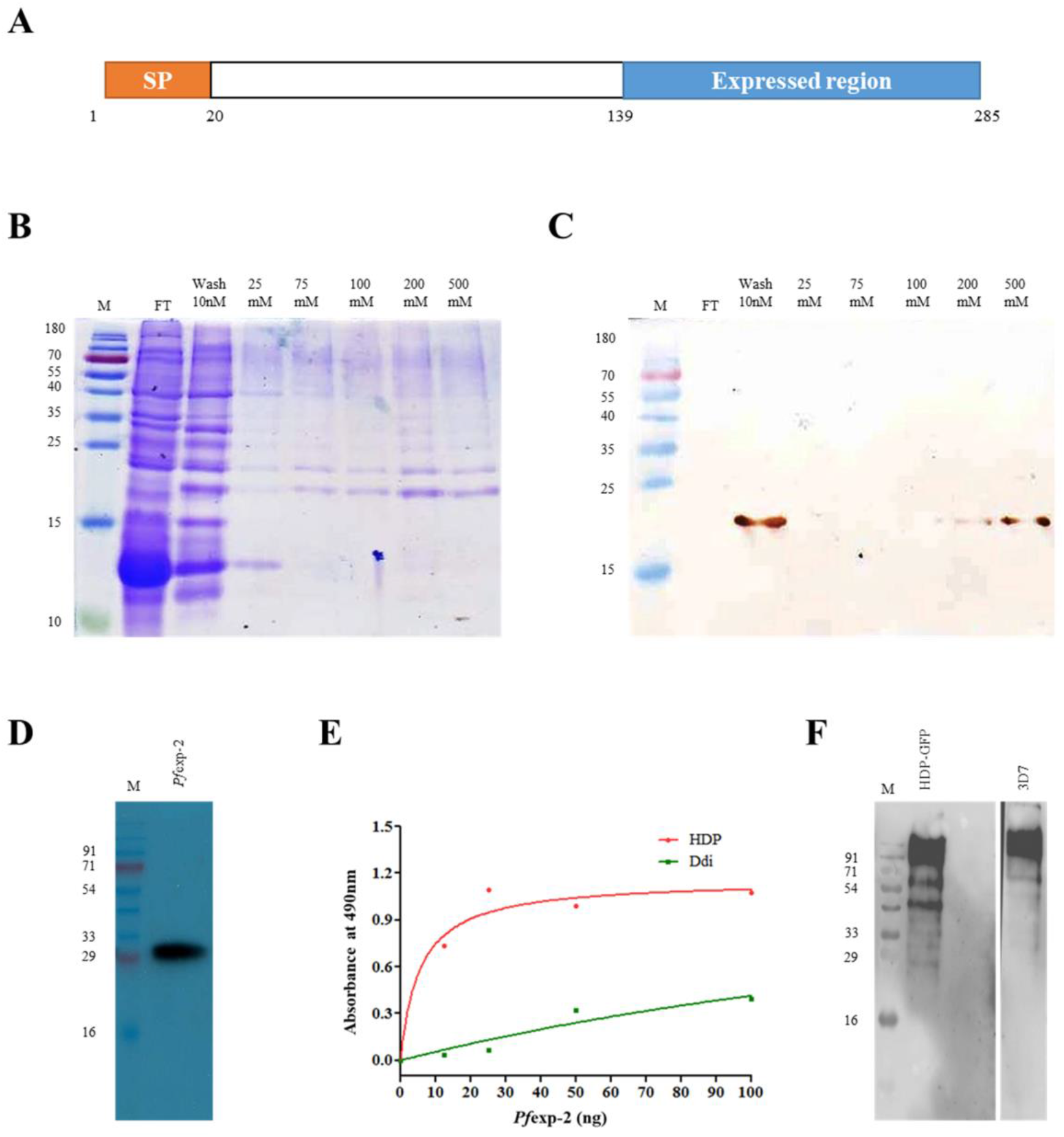
(A) *Pfe*xp-2 - C-terminal from amino acids 139-285 was expressed in pET28b vector in *E. coli*. (B) Coomassie stained SDS-PAGE of different elutes of recombinant *Pfe*xp-2 using urea purification strategy (C) Western blot of different fractions of recombinant purified *Pfe*xp-2 using His-HRP antibody (D) Antibody generated in rabbit against the recombinant *Pf*exp-2 protein recognized the monomeric native protein in 3D7 parasite lysate (E) Recombinant *Pf*exp-2 interacts with *Pf*HDP in a dose dependent manner in an ELISA experiment. Recombinant *Pf*Ddi protein was used as a negative control. (F) western blot analysis of elutes of parasite lysate of *Pf*HDP-GFP parasites immunoprecipitated with GFP antibody using anti-exp-2 antibody detected *Pf*exp-2 in a western blot analysis.

**Sup fig. 5.**
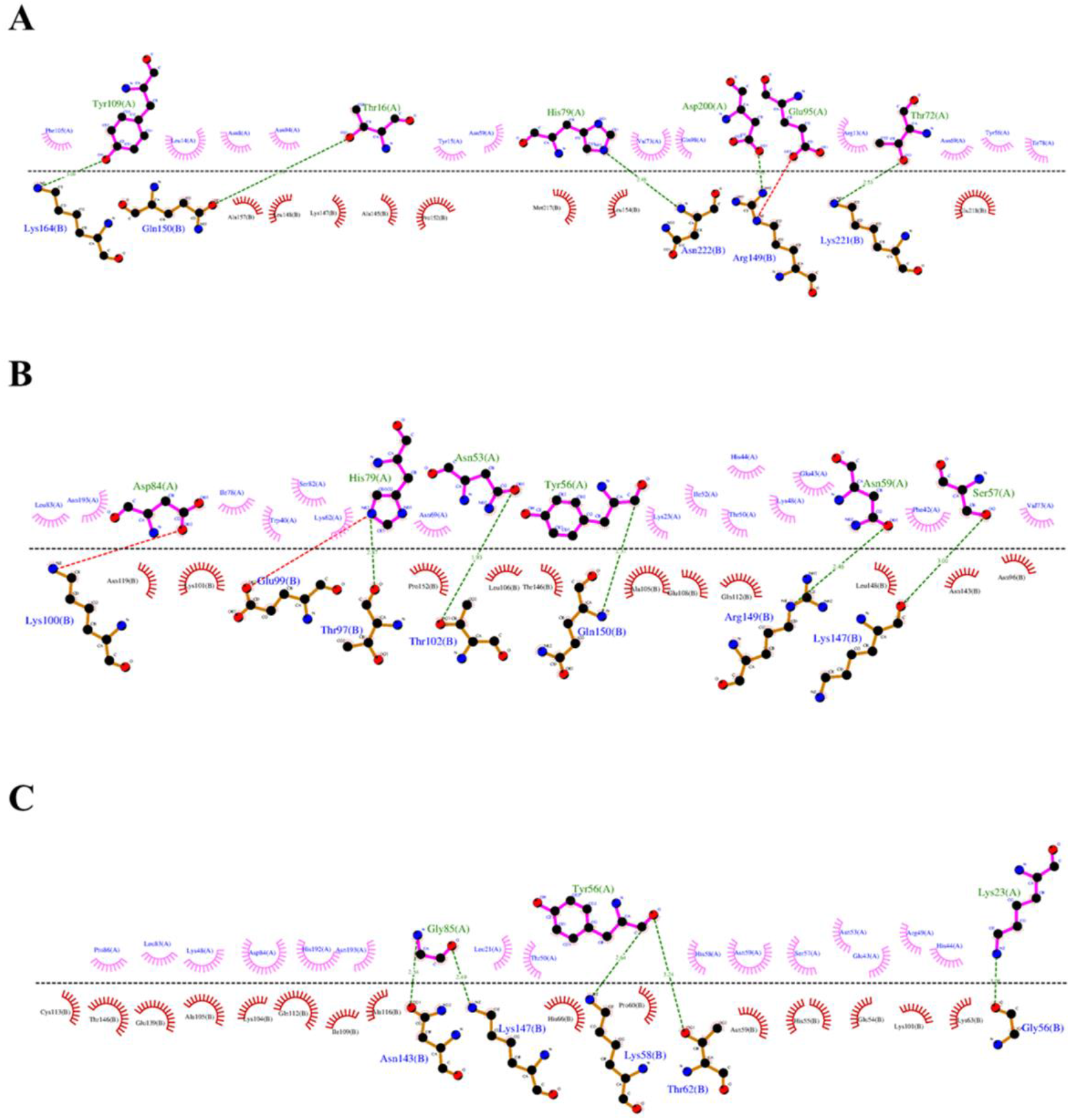
DIMPLOT analysis map for *Pf*exp-2 and *Pf*HDP interacting residues in pose 1 (A), pose 2 (B) and pose 3 (C).

**Sup fig. 6.**
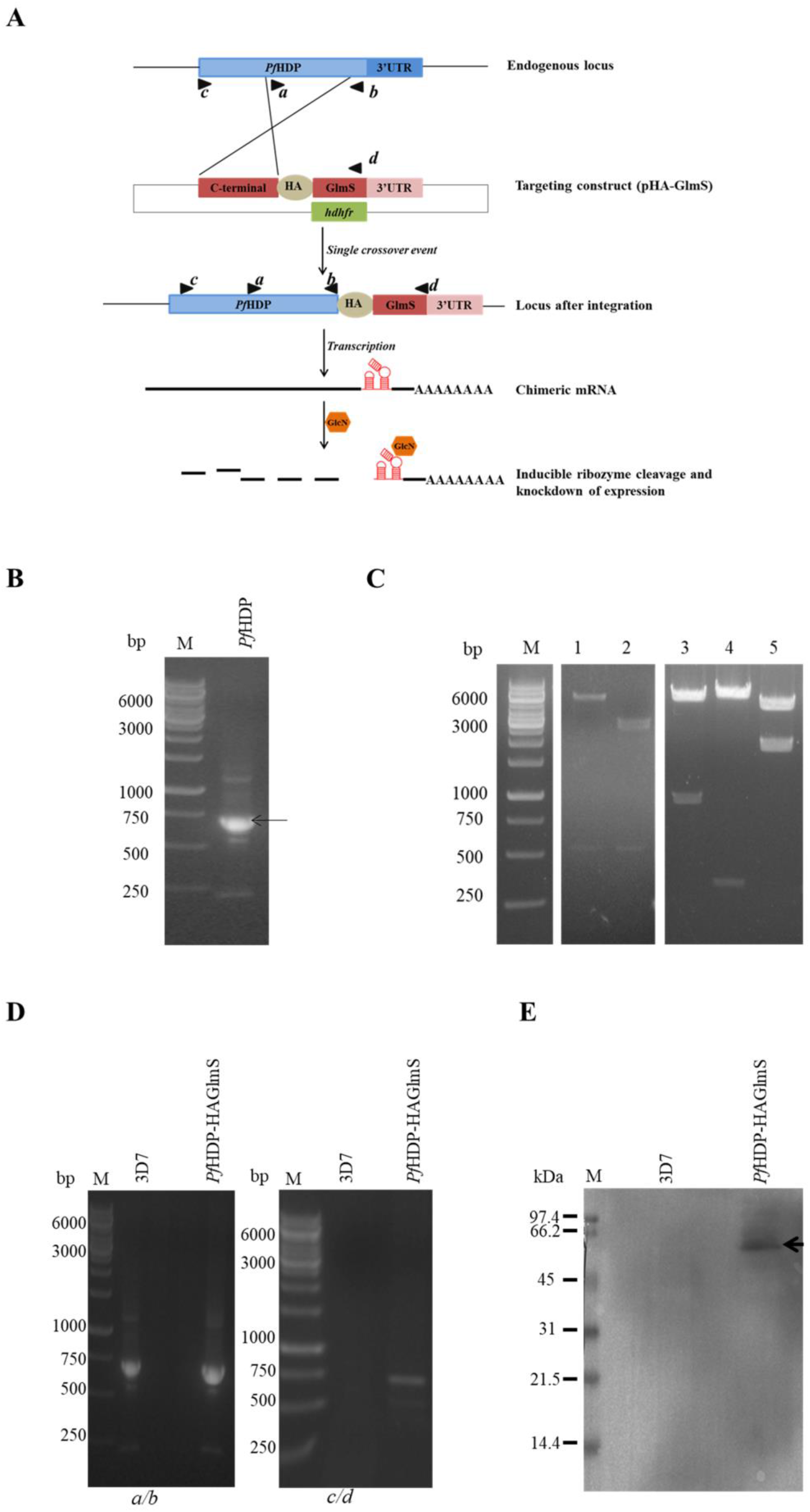
Generation of *Pf*HDP-HAGlmS line in pHA-GlmS vector. (A) Schematic of the GlmS ribozyme reverse genetic tool: The ribozyme is inserted in the 3′-UTR after the coding region so that it is present in the expressed mRNA. Following addition of the inducer, glucosamine, which binds to the ribozyme, the mRNA self-cleaves resulting in degradation of the mRNA and knock down of protein expression. (B) PCR amplification of *Pf*HDP genomic sequence from gDNA using the *Pf*HDPGA_FP and *Pf*HDPGA_RP primer set (C) The construct *Pf*HDP-HAglmS was checked for correct insertion of HDP and presence of other sequence regions using different sets of restriction enzymes (Lane 1: BamHI/HindIII-DHFR; Lane 2: EcoRI/HindIII-HRP; Lane3: BglII/PstI-*Pf*HDP gene, Lane 4: PstI/XhoI-HA_glmS ribozyme, Lane 5: XhoI/SacI: 3′-UTR (D) PCR amplification to check the integration of *Pf*HDP-HAGlmS in parasite genome. (E) Western blot of parasite lysates using anti-HA antibody shows expression of HA tag fused to the *Pf*HDP genomic locus.

**Supplementary Table 1**. The list of *Pf*exp-2 and *Pf*HDP interacting residues involved in protein-protein interaction.

